# A Wnt5a-Cdc42 axis controls aging and rejuvenation of hair-follicle stem cells

**DOI:** 10.1101/2020.10.22.351544

**Authors:** Rajiv L Tiwari, Pratibha Mishra, Nicola Martin, Nikhil Oommen George, Vadim Sakk, Karin Soller, Kodandaramireddy Nalapareddy, Kalpana Nattamai, Karin Scharffetter-Kochanek, Maria Carolina Florian, Hartmut Geiger

## Abstract

Normal hair growth occurs in cycles, comprising growth (anagen), cessation (catagen) and rest (telogen). Upon aging, the initiation of anagen is significantly delayed, which results in impaired hair regeneration. Hair regeneration is driven by hair follicle stem cells (HFSCs). We show here that aged HFSCs present with a decrease in canonical Wnt signaling and a shift towards non-canonical Wnt5a driven signaling which antagonizes canonical Wnt signaling. Elevated expression of Wnt5a in HFSCs upon aging results in elevated activity of the small RhoGTPase Cdc42 as well as a change in the spatial distribution of Cdc42 within HFSCs. Treatment of aged HFSC with a specific pharmacological inhibitor of Cdc42 activity termed CASIN to suppress the aging-associated elevated activity of Cdc42 restored canonical Wnt signaling in aged HFSCs. Treatment of aged mice in vivo with CASIN induced anagen onset and increased the percentage of anagen skin areas. Aging-associated functional deficits of HFSCs are at least in part intrinsic to HFSCs and can be restored by rational pharmacological approaches.

## Introduction

Aging of tissue specific resident stem cells is thought to contribute to tissue attrition in stem cell based tissues both in mice and humans ^1–4^. In the murine skin, aging-associated phenotypic and histological changes include an extended telogen phase of the hair cycle at the expense of anagen as well as miniaturization of hair follicle structures and loss of hair ^5–7^. This loss of hair can be attributed to a decline in the function of hair follicle stem cells (alpha-6 integrin^high^CD34^+^ cells, HFSC) which manifests as a decrease in colony formation activity in-vitro^6^. Mechanistically, the reduced function of HFSCs upon aging has been so far primarily linked to mechanisms extrinsic to HFSCs^8^, like for example reduced responsiveness of aged HFSC to bone morphogenic protein (BMP) and NFATc1 signaling^7^ or proteolysis of Collagen XVII (COL17A1/BP180) which induces terminal differentiation of HFSC towards epidermal keratinocytes^6^. In general though knowledge on HFSC intrinsic aging mechanism are limited, precluding rational approaches to target aging of HFSCs. Wnt signaling is a critical regulator of HFSCs generation and maintenance in young animals^9–12^. In the present study, we demonstrate that stem cell intrinsic non-canonical Wnt5a signaling drives HFSC aging via increasing the activity of the small RhoGTPase Cdc42. Wnt5a can induce a premature aging like phenotype in young HFSCs. Wnt5a inhibition in aged HFSC is able to attenuate aging-associated phenotypes of HFSC. Pharmacological inhibition of Cdc42 activity with the specific inhibitor CASIN enhances canonical Wnt signaling and restores youthfulness of aged HSFC. In-vivo treatment with CASIN induces early onset of anagen and increases hair regrowth in aged mice.

## Results

### A shift from canonical to non-canonical Wnt signaling upon aging in HFSC

Hair growth occurs in a cycle that comprises growth (anagen), cessation (catagen) and rest (telogen). We compared in our analyses primarily hair follicle from day 50 mice and aged mice to identify novel mechanisms of aging of HFSC. While the total number of telogen hair follicle in the back skin of young mice in a “normal” telogen cycle (d-50 mice) ^13,14^ and old mice (>2 years, aged telogen) was similar^6^, some of the telogen hair follicle in aged mice were miniaturized (Fig.S1A-C). We further observed more asynchronized hair growth and a higher percentage of back skin in anagen (characterized by black back skin) in young (day 98, week 14) compared to old mice (Fig.S1D-E). Overall hair follicle structures from the telogen area of young and old skin were though very similar (Fig.S1F, **lower right panel**). Aged animals remained for a longer period in telogen and anagen was consequently delayed compared to anagen duration in young animals (Fig.S1G).

Hair growth in the murine model is driven by a population of stem cells that reside in the bulge and in the hair germ of the hair follicle. These stem cells are termed hair follicle stem cells (HFSCs) with the established markers profile of Sca-1^−/low^alpha-6 integrin (A6)^high^CD34^+^ ^15,16^. Both young and aged animals presented with a similar frequency in HFSCs determined by FACS (Fig.S1H-I) as well as by CD34^+^ immunostaining of tissue sections (Fig.S1J-K). The colony-forming unit (CFU) activity of sorted aged HFSCs was reduced (Fig.S1l), consistent with previously published reports^7^. In summary, HFSC function, but not their number, is reduced upon aging.

Canonical Wnt/ß-catenin signaling plays a critical role for HFSC proliferation and the onset of anagen and thus hair growth in young mice. In the absence of ß-catenin, HFSC fail to induce anagen and fail to produce follicular keratinocytes^9,10,17^. As during aging initiation of anagen was delayed (Fig.S1G), we reasoned that changes in the function of aged HFSCs might be linked to changes in Wnt-signaling^18^. Indeed, expression of canonical Wnt target genes like Axin-2, Lef-1, Lgr-6 and c-Myc was decreased in HFSCs upon aging (Fig.1A and Fig.S1 M). Upon activation of canonical Wnt signaling, ß-catenin stabilizes and translocates to the nucleus to initiate transcription of Wnt target genes. Quantification of nuclear ß-catenin inside the nucleus by 3D quantitative immunofluorescence confocal microscopy demonstrated that the nuclear localization of ß-catenin was reduced in aged HFSC, but not its level of expression (Fig.1B-C and Fig.S1N). Aging of hematopoietic stem cells is for example driven changes in the expression of non-canonical Wnt-ligands within the stem cell^19^. We thus tested expression of Wnt-ligands in young and aged HFSCs. Non-canonical Wnt4 and Wnt7b was decreased while expression of Wnt5a was increased in old HFSC (Fig.1D). While reduced Axin-2 and Wnt4 expression was observed in all compartments of the follicular and interfollicular basal epidermis of aged mice (Fig.S2C-D), increased levels of Wnt5a were restricted to the basal (A-6^high^CD34^+^) and the suprabasal (A-6^low^CD34^+^) compartments of HFSC (Fig.S2E). Among canonical Wnts, Wnt10a presented with elevated levels of expression in HFSCs upon aging (Fig.1D), while the expression of other well characterized ligands like Wnt1, Wnt3, Wnt3a was below our detection level in HFSCs (data not shown).

**Figure 1.**
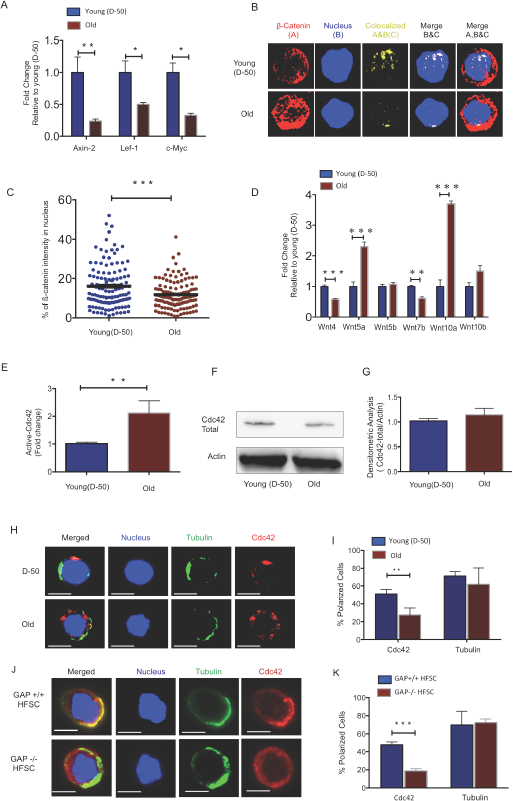
Increased Wnt5a expression in aged HFSC results in a shift from canonical to non-canonical Wnt signaling. **A,** Expression of target genes of the canonical Wnt pathway in FACS sorted young and old HFSC (Sca-1^−/low^A-6^high^CD34^+^), N≥3 **B,** Z-stacks and three-dimensional merged images of ß-catenin (red), nucleus (dapi, blue) and co-localization signal (yellow) in FACS sorted young and old HFSC by Immunofluorescence **C,** Quantification of the ß-catenin signal in the nucleus of young and old HFSC, N=4 **D,** Wnt ligand transcript levels in young (D-50) or old (>2years) HFSC, N≥3 **E,** ICdc42-activity levels in old Sca-1^−/low^ keratinocyte lysate compared to young Sca-1^−/low^ keratinocyte measured by G-LISA, N=4; **F,** Cdc42 protein levels in young and old Sca-1^−/low^ keratinocytes (representative Western blot) **G,** Densitometry score of Cdc42 protein levels from Western blots, N=4 **H,** Representative picture of Cdc42 (red) and tubulin (green) in young and old HFSC, scale bar = 5μm **I,** Percentage of cells with a polar distribution of Cdc42 and tubulin in young and old HFSC, N≥6 **J,** Representative picture of a non-polar distribution of Cdc42 (red) in young HFSC from Cdc42GAP^−/−^ as compared to HFSC from young wild type Cdc42GAP^+/+^ mice, scale bar=5μm **K,** Quantification of polar distribution (percentage) of Cdc42 and tubulin in HFSC from Cdc42GAP^+/+^ and Cdc42GAP^−/−^ mice, N≥3, P<0.05, **P<0.01, ***P<0.001 (paired student’s t test). Error bars represent s.e.m.

Cell division cycle 42 (Cdc42) is a small GTPase of the Rho family. It cycles between two conformational states; an active, GTP bound and an inactive, GDP bound state^20^. Like all GTPases, in the active state, Cdc42 can bind distinct effector proteins to then cell-type specifically activate distinct types of signaling pathways^21^. We previously reported an about 2-fold increase in the activity of Cdc42 in primitive aged hematopoietic cells ^22,23^. Indeed, Cdc42 activity was also about 2-fold elevated in an cell population enriched for aged HFSC (Sca-^1-/low^ cells) (Fig.1E), while the level of Cdc42 itself remained unchanged (Fig.1F-G and Fig.S1O). Increased Cdc42 activity induces an apolar distribution of Cdc42 itself as well as of tubulin and other polarity proteins in aged HSCs^22,24^. Similar to aged HSCs, aged HFSCs presented with a decrease in the percentage of cells polar for the distribution of Cdc42, but unlike aged HSCs, aged HFSCs did not show an apolar distribution of tubulin (Fig.1H-I). We also investigated, in addition to Cdc42 and tubulin, the distribution of Par6 and Numb that are also polarity proteins and known to mark the apicobasal axis of epithelial cells. Par6 showed a polar distribution in both young and aged HFSC, while the frequency of HFSCs polar for Numb decreased upon aging (Fig.S2D, G-I). The Cdc42GAP protein is a negative regulator of Cdc42 activity. Cdc42GAP^−/−^ mice present therefore with a constitutive increase in the activity of Cdc42 in all tissues^25^ and display a premature aging like phenotype which also includes reduced hair regeneration^25^. HFSC from young Cdc42GAP^−/−^ mice were apolar for the distribution of Cdc42 (Fig.1J-K), demonstrating that apolarity of aged HFSC is tightly linked to an increased Cdc42 activity in HFSCs, similar to what has been previously described for aged HSCs^22^.

### Wnt5a induces aging of HFSCs

Wnt5a, expressed within primitive hematopoietic cells or given exogenously, increases the activity of Cdc42 in primitive hematopoietic cells^23,19^. Aged HFSC show elevated levels of expression of Wnt5a (Fig.1D). We thus tested first whether indeed Wnt5a is also able to increase Cdc42 activity in HFSCs. Treatment of young HFSCs with Wnt5a (300ng/ml) increased the activity of Cdc42 to a level seen in aged HFSCs, without affecting the level of Cdc42 itself (Fig.2A-C). Young HFSCs treated with Wnt5a presented with a reduced frequency of cells polar for Cdc42 (Fig.2D-E), which is consistent with a role of elevated Cdc42 activity in reducing the frequency of HSCs polar for the distribution of polarity proteins^19^. Non-canonical Wnts usually antagonizes canonical Wnt signaling ^26,27^. We therefore tested if Wnt5a inhibits canonical Wnt signaling in HFSC. Treatment of young HFSCs with Wnt5a decreased the amount of ß-catenin with a nuclear localization and the expression of the canonical Wnt target gene Axin-2 (Fig.2F-H). Wnt5a treatment further reduced the colony forming activity of young HFSC (Fig.2I-J). We then reduced Wnt5a expression in aged Sca-^1-/low^ keratinocytes by a lentiviral knockdown approach (Fig.S3A-C). Reduced expression of Wnt5a in aged cells re-induced Axin-2 expression (Fig.S3D) and increased the colony forming activity of aged Sca-1-/low keratinocytes (Fig.2K-L and Fig.S3E). These data show that non-canonical Wnt5a-Cdc42 signaling antagonizes canonical-Wnt signaling in aged HFSCs. These data also demonstrate that the aging-associated decline in CFU activity can be restored to a youthful level by reducing the level of expression of Wnt5a in aged HFSCs. This implies that the cell-intrinsic increase of Wnt5a expression in aged HFSC causatively contributes to the decline in function of aged HFSCs.

**Figure 2.**
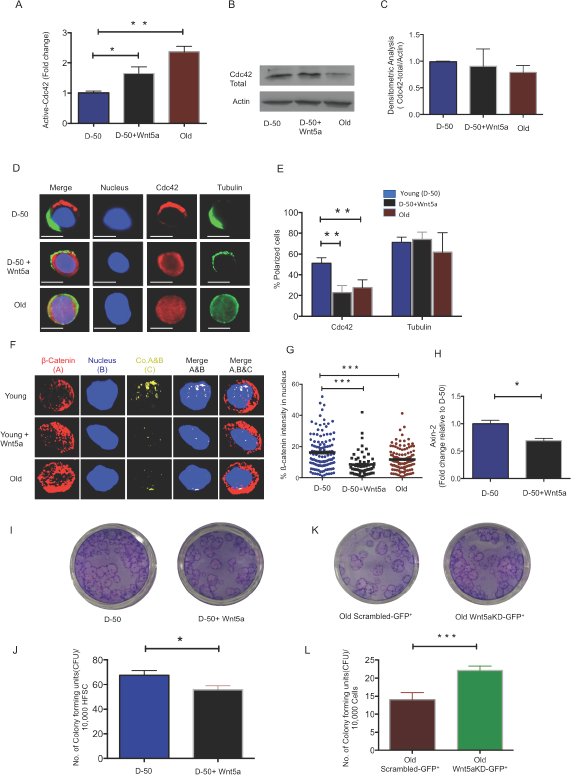
Wnt5a regulates Cdc42 activity and induces an old like phenotype in young HFSCs. **A,** Cdc42 activity measured using G-LISA kit in lysate of young Sca-1^−/low^ keratinocyte after Wnt5a treatment for 2 hours, N=4 **B**, Representative Western blot on total Cdc42 protein levels in Wnt5a treated young Sca-1^−/low^ keratinocytes **C,** densitometric score of western blot results for total Cdc42 in young Sca-1^−/low^ keratinocyte after 2 hours of Wnt5a treatment N=3 **D,** Cdc42 (red) and tubulin (green) distribution in young and old HFSC with or without Wnt5a treatment for 2 hours scale, immunofluorescence, scale bar=5μm **E,** Percentage of cells polar for the distribution of Cdc42 and tubulin in young and old HFSC after Wnt5a treatment for 2 hours, N≥3 **F,** Z-stacks and three-dimensional merged images of ß-catenin(red), nucleus (dapi, blue) and their co-localization (yellow) in FACS sorted young and old HFSC by immunofluoresence **G,** Quantification of the ß-catenin signal in the nucleus and cytoplasm of young, old and Wnt5a treated HFSCs, N=3 **H,** Expression levels of Axin-2 in young HFSC after Wnt5a treatment, N=3 **I,** Morphology of colonies formed by young HFSCs after Wnt5a treatment **J,** Number of colony forming units in young, aged and Wnt5a treated HFSCs, N=3 **K,** Morphology of colonies formed by old Sca-1-/low keratinocytes transduced (green) with non-targeted scrambled (NT-GFP+) and targeted Wnt5a knock down (Wnt5aKD-GFP+) sh-RNA **L,** Number of colony forming units in old Sca-1-/low keratinocyte transduced (green) with non-targeted scrambled (NT-GFP+) and targeted Wnt5a knock-down (Wnt5aKD-GFP+) sh-RNA, N=3, *P<0.05, **P<0.01, ***P<0.001 (paired student’s t test). Error bars represent s.e.m.

**Figure 3.**
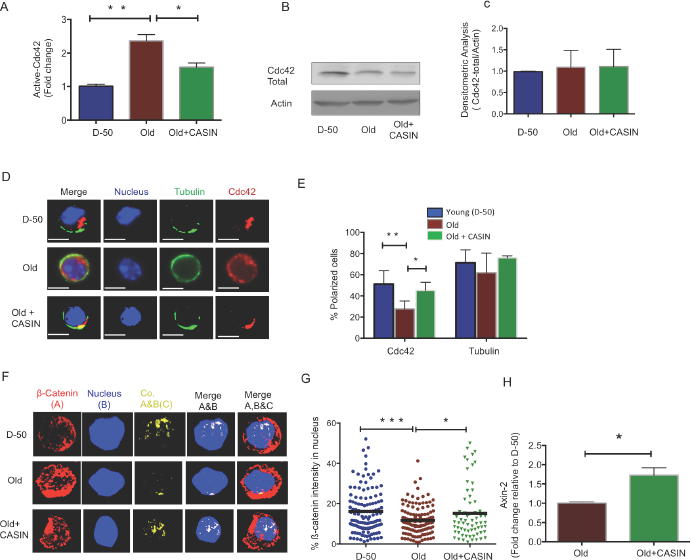
Inhibition of Cdc42 activity with CASIN induces a young like phenotype in old HFSCs and re-establishes canonical Wnt Signaling. **A,** Cdc42 activity measured by a G-LISA in lysate of old Sca-1^−/low^ keratinocyte cultured for 2 hours with the Cdc42 activity inhibitor CASIN, N=4 **B,** Representative Cdc42 protein level in young, old and CASIN treated old Sca-1^−/low^ keratinocytes by Western Blot **C,** Densitometric score for the amount of Cdc42 protein in Sca-1^−/low^ keratinocyte from young, old and old CASIN treated for 2 hours, N=3 **D,** Distribution of Cdc42 (red) and tubulin (green) in young, old and old HFSCs cultured for 2 hours with CASIN by immunofluoresence, scale bar=5μm **E,** Percentage of cells with a polar distribution of Cdc42 and tubulin in young, old and CASIN treated old HFSC cultured for 2hrs, N≥3, **F,** Z-stacks and three-dimensional merged images of ß-catenin (red), nucleus (DAPI, blue) and their co-localization (yellow) in FACS sorted young, old and CASIN treated HFSC cultured for 2 hours by immunofluorescence **G,** Quantification of the amount of nuclear ß-catenin in young, old and CASIN treated old HFSC cultured for 2 hours, N=3, **H,** Level of expression of Wnt target genes in old HFSC cultured with CASIN for 6 hours, N=3, *P<0.05, **P<0.01, ***P<0.001 (paired student’s t test). Error bars represent s.e.m.

### Pharmacological inhibition of Cdc42 promotes anagen and hair regrowth in old mice in vivo

Elevated activity of Cdc42 in aged HSCs causes aging of HSCs^22,28^. We thus tested whether elevated levels of activity of Cdc42 seen in HFSCs might be also critical for aging of HFSCs. The activity of Cdc42 can be specifically inhibited by CASIN (Cdc42 activity specific inhibitor)^29,30^. Treatment of aged Sca-^1-/low^ keratinocytes with 10μM of CASIN resulted in a reduction of Cdc42 activity to the level seen in young HFSCs (Fig.3A), without affecting the level of total Cdc42 (Fig.3B-C). Aged HFSCs treated with CASIN presented with a youthful percentage of HFSCs polar for the distribution of Cdc42 as well as Numb (Fig.4D-E and Fig.S4A-B). Aged CASIN treated HFSCs also presented with an increase in the nuclear localization of ß-catenin and with elevated expression of Axin-2 (Fig.3F-H). A youthful level of Cdc42 activity in chronologically aged HFSCs restores a youthful level of cell polarity and canonical Wnt signaling in aged HFSCs.

**Figure 4.**
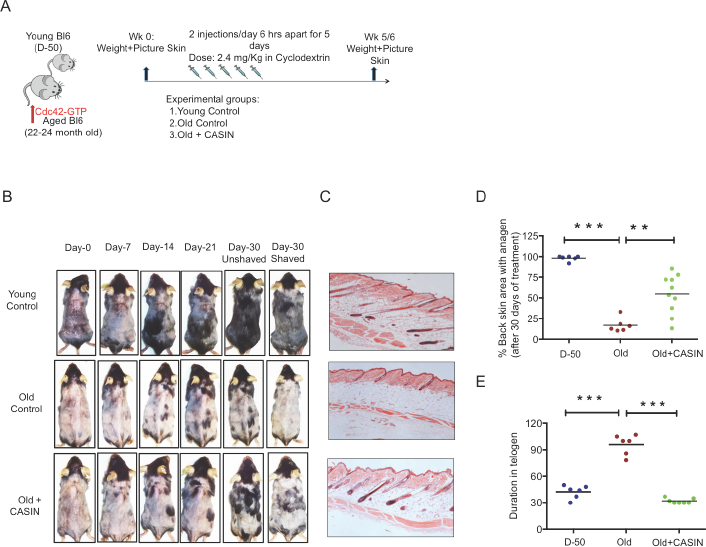
In vivo treatment with CASIN induces onset of anagen in aged mice. **A,** Experimental setup **B**, Back skin of representative aged mice and aged mice treated i.p with CASIN, pictures were taken weekly **C**, H&E staining of mouse back skin sections (longitudinal) from young, aged animals and the anagen area of skin from aged animals after CASIN treatment **D**, Quantification of the size of the skin area with anagen in D-50, old and old CASIN treated animals N≥6 **E**, Duration of telogen in young, old and old CASIN treated, N≥6, *P<.05, **P<.01, ***P<.001 (paired student’s t test). Error bars represent s.e.m.

**Figure 5.**
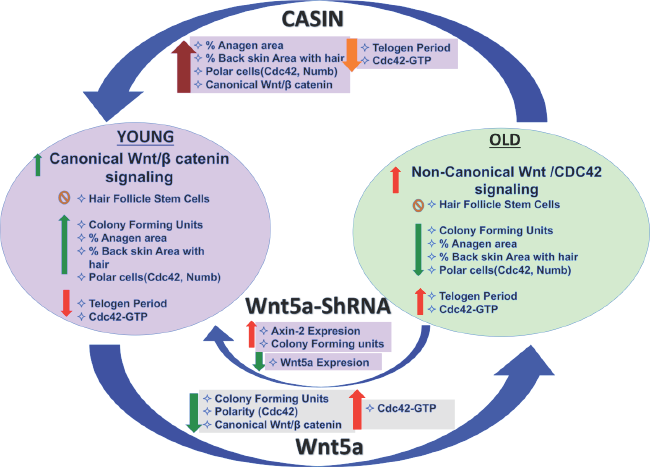
Graphical summary

Canonical Wnt signaling is critical for the onset of anagen in HFSCs (24). As inhibition of Cdc42 activity via CASIN was able to increase canonical Wnt signaling in aged HFSCs in vitro, we finally tested whether inhibition of Cdc42 activity via CASIN in vivo ^9,11,29^ might also restore anagen onset in aged skin in vivo. To this end, aged mice were given CASIN twice a day for 5 days (Fig.4A). The level of hair growth was subsequently determined for up to 30 days post treatment (Fig.4B). Anagen onset in CASIN treated aged animals was analyzed by HE staining of the black area of the back skin (Fig.4C). Anagen areas (black patches) were about 3-fold more frequent in CASIN treated aged mice in comparison to untreated controls (Fig.4D). In addition, the duration of telogen in aged CASIN treated animals was reset to the duration of telogen in young (D-50) mice (Fig.4E). In summary, CASIN, when provided in vivo, enhanced anagen onset in the skin of aged mice to allow for growth of novel hair.

## Discussion

We report an increase in the activity of the small RhoGTPase Cdc42 in aged HFSCs, driven by an intrinsic increase in Wnt5a in HFSCs. This negatively impacts HFSC function, likely by antagonizing canonical-Wnt signaling. Overexpression of Wnt5a in mice has been reported to cause an extended telogen period, a decrease in beta catenin/TCF reporter activity and loss of hair due to inhibiting of the second and third wave of hair follicle formation ^31–33^, which are phenotypes consistent with aging. Our data do not exclude that Wnt5a secretion from niche cells other than differentiated and undifferentiated keratinocytes (Fig.S2E) might also influence HFSC aging, though Wnt signaling usually acts on a very short range ^34^. Consistent with Lim et al^12^ and in contrast to earlier reports^15,35,36^ our data support a persistent activation of canonical Wnt-signaling in HFSC throughout both telogen and anagen in HFSC from young (D-50) as well as old (>2years) mice, as indicated by the expression of Axin-2. However, levels of activation are overall lower upon aging (Fig.1A-C).

Inhibition of Wnt5a or pharmacological targeting of Cdc42 activity by CASIN in aged HSFCs restores a youthful function of HFSCs both in vitro. A critical role of elevated activity of Cdc42 in stem cell aging^19,37^ is also consistent with the already published impaired hair regeneration phenotype of Cdc42GAP^−/−^ mice ^22,25^. Cdc42GAP^−/−^ mice present with constitutively increased Cdc42 activity already in young animals. CASIN re-establishes a youthful level of canonical Wnt-signaling in chronologically aged HFSCs including elevated Axin-2 expression and nuclear beta catenin *in-vitro* (Fig.3F-H). CASIN also induces anagen onset in aged mice *in vivo* (Fig. 4E). We recently demonstrated that inhibition of Cdc42 activity in aged mice in vivo extends the average and maximum life span ^29^. For extension of life-span, CASIN was administered once for 4 days in a row instead of 5 days twice for experiments reported here, and provided at a higher overall dosage (2.4mg/kg per injection here vs. 25 mg/kg for the extension of lifespan). Distinct ways of administration of CASIN and levels in vivo might thus be able to elicit distinct positive biological effects in vivo^29^.

Elevated levels of Wnt5a and increased Cdc42 activity alter HSFCs polarity. A role for Wnt5a and Cdc42 in controlling changes in cell polarity in other cell types have been previously reported^19,22,37^. A polar phenotype of Cdc42 in HSCs is associated with a strong regenerative capacity, similar to what is here reported for HFSCs ^38^. Cdc42 itself is part of the cytoplasmic protein polarity complex (Par6-Cdc42-aPKC) that is known to regulate the activity of PKC to maintain apico-basal polarity in epithelial cells ^39,40^. Interestingly, genetic loss of atypical aPKCλ has been shown to alter the fate determination of HFSC^41,42^ with a phenotype similar to that of aged HFSCs. In summary, our data demonstrate that intrinsic aging of HFSCs is linked to an elevated activation of the Wnt5a-Cdc42 axis that can be attenuated by targeting Cdc42 activity in vivo.

Finally, whether the number of HFSCs change with age still remain controversially discussed ^6,7,43^, while there is consensus that the function of aged HFSCs is reduced. For example Matsumura et al^6^ reported a decrease in HFSC number in the hairless region of the back skin of aged mice, while our data support that the number of HFSCs is not altered upon aging. Further studies will thus be necessary to unequivocally address the level of change in the number of HFSCs upon aging.

## Materials and Methods

### Animals

C57BL/6 mice were obtained from the animal facility, Ulm University, Germany. Mice were used as D-50 (Week-8 as young) or >2years for old until otherwise noted. Cdc42GAP−/−mice were obtained from the animal facility at CCHMC, Ohio, USA. Animals were housed and handled in accordance with the IACUC at CCHMC and the Regierungspräsidium Tübingen permission numbers 0.165, 1172 and 1296 respectively.

### Reagents

The antibodies to cdc42, Numb goat polyclonal antibody, PAR6 and Tubulin were obtained from Abcam. The Alexa Fluor 488 (Donkey Anti-Rabbit and Anti-Rat IgG H&L) were also obtained from Abcam. The APC-alpha-6Itg anti human/mouse CD49f was obtained from BioLegend^R^, Biotin anti-mouse Sca-1 (Ly-6 A/E) antibody (Clone D7) was obtained from eBioscience. Cy3^R^ (Donkey Anti-Rabbit and Anti-Goat IgG H&L was obtained from Abcam). Dynabeads Sheep anti-rat igG was obtained from Invitrogen, life technologies. FcR block anti mouse CD16/CD32 and PE cy7 antibody was obtained from eBiosciences. PE-CD34 Rat anti-mouse CD34 was obtained by BD Pharmigen. Streptavidin-PEcy7 was obtained from eBioscience. Sytox blue was obtained from ThermoFisher Scientific.

### Determination of the size of anagen area

Black patches onto the mice skin are considered to be in anagen ^7^. To measure the percentage of mouse back skin with anagen area, mice skin was shaved and photographed. The percentage of area of mouse back skin with black spots was calculated with Imaging processing and quantification software (Adobe Potoshop) using a square of 4 cm^2^ area as control for intensity.

### Paraffin sections and H&E staining

For paraffin section mouse back skin was fixed in 4% formalin solution overnight at 4°C and was embedded in the paraffin. Paraffin embedded skin section were cut into 5-μM-thick sections and used for the staining with H&E and IF staining. For H&E staining paraffin section were deparaffinized and then rehydrated and stained with hematoxylin and eosin.

### FACS analyses and isolation of HFSC

FACS isolation of HFSC was performed using published protocols ^44,45^. Briefly back skin of the mice was cut with scissors and kept in ice cold PBS. With the help of scalpel subcutaneous fat was removed and skin was kept in Dispase solution (4μg/ml), dermal side down and incubated overnight at 4°C. Epidermis was peeled out and treated with trypsin (.025%) in HBSS for 10 min at 37°C to get a single cell suspension. The stem cell fraction was enriched by magnetic depletion using biotinylated Sca-1Ab and the Dynabead system. After magnetic depletion cells were stained with anti-alpha-6 integrin (Biolegends, clone GoH3) and anti-CD34 (clone RAM) (eBioscience).

### Colony formation assay

Colony formation assay was performed on sorted HFSCs (Sca-1^−/low^A-6^high^CD34^+^). For this 10,000 HFSC were plated in lethally irradiated 3T3 feeder cells in Ca-Mg-Free FAD basal medium containing EGF, Insulin, hydrocortisone supplemented with Ca-free FBS for 14 days and medium was changed every second day. On day 14 cells feeder layer was removed with Trypsine:Versene (1:5) solution followed by fixing in 4%PFA for 20 min and staining in 1% of Rhodamine and 1% of Nile blue in water^45^.

### Immunofluorescence staining of paraffin sections

Paraffin section was used for immunofluorescence staining of CD34(1:200), Wnt5a (1:100) and active Cdc42(1:50),(EMD Millipore)^46^. Antigen retrieval was performed for both the antigen by boiling the paraffin sections after deparaffinization in Dako target retrieval buffer for 20 minutes (Dako, Carpentaria, CA, USA). Sections were blocked for nonspecific binding in BSA in PBS containing 10% goat serum. Primary Ab incubation was performed for overnight at 4°C followed by fluorescence conjugated secondary antibody incubation. DAPI was used to counterstain the nucleus. Stained samples were analyzed on a fluorescence or a confocal microscope.

### Immunofluorescence staining of sorted HFSC for polarity analysis

Freshly sorted HFSC were seeded on fibronectin coated glass coverslips. For polarity analysis, HFSC were incubated for 2 hrs with 300 ng/ml Wnt5a and 10μM CASIN or left untreated. After incubation at 37°C, 5% C0_2_ and 3% O_2_ in growth factor free medium, cells were fixed with BD Cytofix Fixation Buffer (BD Biosciences). After fixation cells were gently washed with PBS and permeabilized with 0.2% TritronX-100 (Sigma) in PBS for 20 min and blocked with 10% donkey serum for 30 min. Primary Ab incubation followed with secondary antibody incubation were performed at room temperature for 1hrs. The coverslip was mounted with ProLong Gold Antifade reagent with DAPI (Invitrogen, Molecular Probes). Cells were stained with an anti-Cdc42 (Millipore, rabbit polyclonal), an anti-ß-catenin (Millipore, rabbit polyclonal), Par6 (Santa Cruz Biotechnology, goat polyclonal), Tubulin (rat, monoclonal), followed by incubation with secondary antibodies conjugated with Alexa fluor 488 and Cy-5. Samples were imaged with an AxioObserver Z1 microscope (Zeiss) with a X63 objective. Images were analyzed with Axio Vision 4.6 software. Polarity scoring was performed based on the localization of each single stained protein, if it was asymmetrically distributed with respect to a plane through middle of the cell. Alternatively, samples were analyzed with LSM710 confocal microscope (Zeiss) equipped with a X63 objective. Primary raw data were imported into the Velocity Software package (Version 6.0, Perkin Elmer) for further processing and conversion into three-dimensional images. Analysis of the localization analysis of ß-catenin was performed using velocity software (percentage of ß-catenin intensity in the nucleus above the threshold level).

### G-LISA

For the determination of active GTP bound form of Cdc42 we used the G-LISA kit for Cdc42 from Cytoskeleton according to the protocol of the manufacturer.

### Reverse-transcriptase real time PCR

20,000-40,000 HFSC from young and aged mice were lysed and processed for RNA extraction immediately after sorting or after treatment of CASIN and Wnt-5a for 2hrs. RNA was obtained with microRNA extraction kit(Qiagen) and whole RNA was used for cDNA preparation. cDNA was prepared and amplified with Ovation RNA amplification system V2 (Nugen). All real-time PCR reaction was performed using Taqman real time PCR reagent and primers from Applied Biosystem on an ABI9700HT real time PCR machine.

### Western blot

For the measurement of protein expression, western blot was performed on Sca-1^−/low^ keratinocyte or in keratinocyte cell lysate from back skin of young and old mice for Cdc42 using anti-Cdc42 (Millipore, rabbit polyclonal) antibody and was normalized with the actin blot using the Actin (Sigma) antibody. The relative level of expression was estimated by densitometry quantification.

### Lentivirus mediated knockdown of Wnt5a

Aged mice (24-month-old) were killed; Sca1-low cells were isolated from the back-skin area as previously described. These cells were further transduced overnight on retronectin-coated (TaKaRa) plates with cell-free supernatants containing lentiviral particles according to reference^19,47^. The lentivirus plasmid vector pLKO 1-YFP was obtained from Sigma’s validated genome-wide TRC shRNA libraries (Sigma-Aldrich) and was further changed to eGFP in-house.

### Statistical analysis

A minimum 3 and up to 6-7 replicates was done for experiments presented. Data are presented as mean and standard error means (SEM). Comparison between groups has been done using Student’s t-test assuming two tailed distribution and unequal variances. Differences were considered statistically significant at the p<0.05 level.

## Acknowledgments

We thank Heidi Hainzl, Karmveer Singh and Pallab Maity for advice and critical support for initiating some of the experiments. We thank the flow core at Ulm University and at CCHMC for their support, and A. Rück and J. Breymayer from the imaging core at Ulm University for support with confocal microscopy, and the Mouse and Cancer Core in Cincinnati and the Tierforschungszentrum of the University of Ulm for supporting our animal work. The work in the laboratory of H.G. was supported by grants from the German Federal Ministry of Education and Research (BMBF) within its joint research project SyStaR (also to KSK) and the Excellence program of the Baden-Württemberg Foundation.

## Author Contribution

RLT and HG contributed to study conception, experiment design, data acquisition and analysis and manuscript writing, PM and NOG helped in data acquisition and analysis, NM,VS,KSD,KN helped in data acquisition, KRN, CMF and KSK contributed to manuscript writing and critical revision.

---

**Figure S1.**
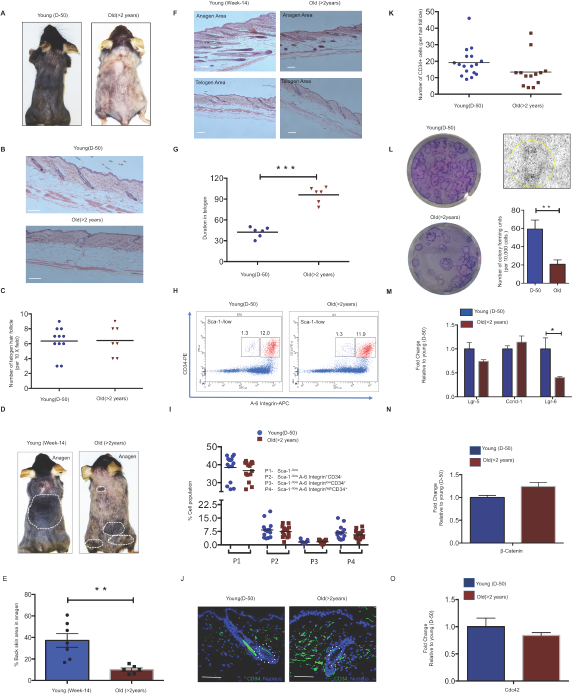
Functional decline of aged HFSC. **A,** Young (D-50) and old (>2years) mice in telogen stage, hair shaved, **B,** H&E staining of a longitudinal section of back skin from young (D-50) and old (>2years) mice (miniaturized hair follicle), scale bar = 100μm **C,** Number of telogen hair follicle counted per 10X field picture of telogen mice skin of young D-50 and old >2 years, N≥7 **D,** Morphology of young (week-14) and old(>2years) mice in anagen stage after hair shaving, anagen area are encircled by white lines **E**, measurement of anagen areas from young (week-14) and old(>2years) mice and represented as percentage of complete back skin area, N≥6 **F,** Representative picture of H&E stained LS of back skin from, anagen area(upper panel), telogen area (lower panel) of young (week-14) and old(>2years) mice, yellow yarrow shows hyper proliferated bulge morphology, Scale bar = 100μm **G,** Duration of telogen in young(D-50) and old mice(>2years) showing increase in duration of old HFSC in telogen stage, N=6 **H**, FACS analysis of Sca-1-/low fraction to detect different cell population in young and old mice **I,** Percentage of different cell population in epidermal cell suspension of skin from D-50 and Old mice showing no change in HFSC number, N≥14 **J,** Immunofluorescence images with HFSC marker CD34 (green and nucleus (blue) in LS of telogen skin from young(D-50) and old mice **N**≥3, Scale bar = 50μm **K,** Quantification of number of CD34+ cells counted per telogen hair follicle in young(D-50) and old (>2years) telogen hair follicle, **L,** Representative picture of colony forming units from FACS sorted HFSC from young and old mice in telogen(left panel),Representative morphology of colonies obtained during CFU analysis (right upper panel), Quantification of CFU per 10,000 HFSC plated from young and old mice indicating decrease in colony forming units in old HFSC (right lower panel), **M,** Transcript levels of canonical Wnt target genes (Lgr-5, Ccnd-1 and Lgr-6) **N**, ß-catenin **O,** Cdc42 in young and old HFSC **N**≥3,*P<.05, **P<.01, ***P<.001 (paired student’s t test). Error bars represent s.e.m.

**Figure S2.**
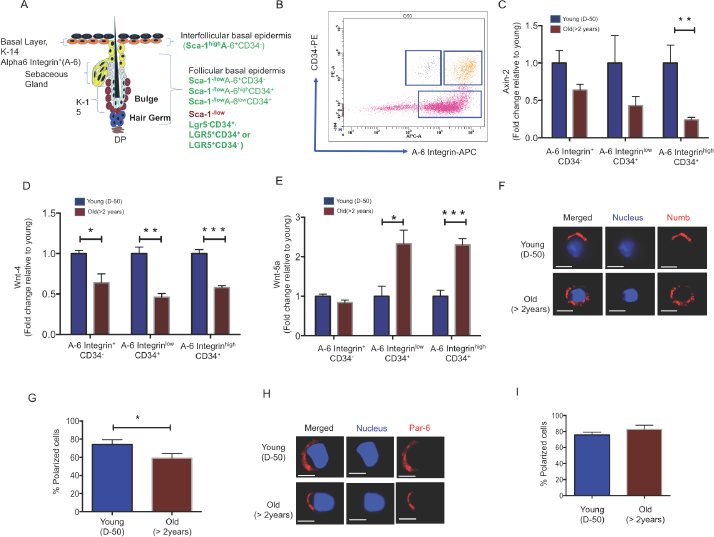
Increase in Wnt5a expression upon aging is specific to a CD34^+^ cell population. **A**, Graphical representation of the structure of the hair follicle and location of distinct types of stem and progenitor cells **B**, dot plot of FACS analyses on Sca-1^−/low^ keratinocytes showing three different cell (A-6^+^CD34^-^, A-6^low^CD34^+^ and A-6 ^high^ CD34^+^) populations obtained during FACS sorting **C**, transcript levels of Axin-2 **D**, Wnt4 and **E**, Wnt5a in young and old A-6^+^CD34^-^,A-6^low^CD34^+^ and A-6 ^high^ CD34^+^ cells, N≥3 **F,** Representative immunofluorescence image 10X and **G,** 40X of Wnt-5a (red) CD34(green) and Nucleus (dapi, blue) in telogen hair follicles, Polarity of Numb and Par-6 in young and old HFSC **H,** Representative picture of the distribution of Numb (red) in young and old HFSC, immunofluorescence, Scale bar =5μm **I,** Percentage of cells polar for Numb in young and old HFSC, N≥3 **J,** Distribution of Par-6 (red) in young and old HFSC, immunofluorescence, Scale bar =5μm **K,** Percentage of cells polar for Par-6 in young and old HFSC, N≥4, *P<.05, **P<.01, ***P<.001. (paired student’s t test). Error bars represent s.e.m.

**Figure S3.**
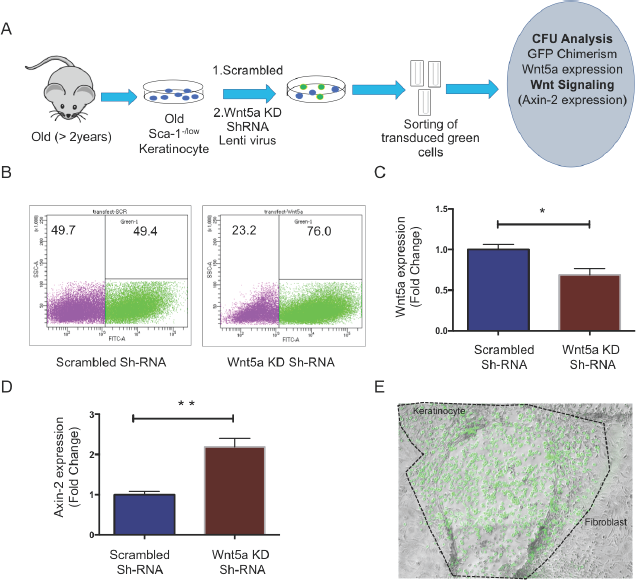
Suppression of expression of Wnt5a in old HFSC induces a young-like phenotype. **A,** Experimental setup **B,** Representative FACS plot showing percentage of transduced cell in scrambled shRNA and Wnt5a knock-down shRNA transduced cells on day 7 post transduction, N≥3 **C,** level of expression of Wnt5a in scrambled shRNA and Wnt5a knock-down shRNA transduced cells on day 7 post transduction, N≥3 **D,** Expression of canonical Wnt-target genes in scrambled shRNA and Wnt5a knock-down shRNA transduced cells on day 7 post transduction, N≥3 **E,** Representative picture of green colony formed by a transduced old Sca-1-/low keratinocyte, surrounding fibroblast feeder cells are non-green, **N**≥3. *P<.05, **P<.01 (paired student’s t test). Error bars represent s.e.m.

**Figure S4.**
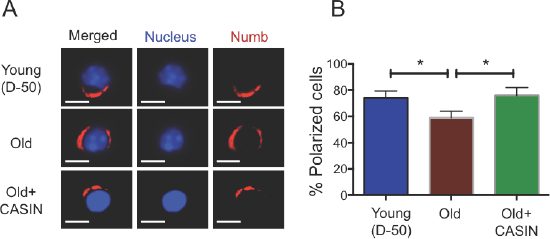
CASIN treatment of aged HFSCs restores a youthful level of cell polar for Numb. **A,** Representative picture of Numb (red) in young, old and CASIN treated HFSC, Immunofluorescence, N≥3 **B,** Percentage of cells polar for Numb in young, old and old CASIN treated HFSCs, N≥3. Scale bar =5μm, *P<.05. (paired student’s t test). Error bars represent s.e.m

